# Identification of Novel Cryptic and Classical Clades in *Clostridioides difficile*

**DOI:** 10.1101/2025.08.01.668003

**Authors:** Enzo Guerrero-Araya, Francisca Cid-Rojas, Marina Muñoz, César Rodríguez, Daniel Paredes-Sabja

## Abstract

*Clostridioides difficile* is a major cause of nosocomial infections and antimicrobial-associated diarrhea, with significant global health implications. While five classical clades (C1 to C5) have traditionally encompassed most pathogenic strains, recent genomic studies have uncovered several highly divergent cryptic clades (C-I to C-V), suggesting greater taxonomic and evolutionary complexity. In this study, we performed a comprehensive whole-genome analysis of 25,144 publicly available *C. difficile* genomes, supplemented with 21 novel isolates from Costa Rica and Brazil. Using average nucleotide identity (ANI), recombination-corrected core-genome phylogenies, we confirmed the presence of all known classical and cryptic clades and identified two novel classical clades (C6 and C7) and two previously undescribed cryptic clades (C-VI and C-VII). We further detected evidence for bifurcation within cryptic clade C-III, supporting its division into two lineages C-IIIa and C-IIIb. ANI comparisons revealed that cryptic clades share less than 95% identity with classical C. difficile strains but remain more similar to *C. difficile* than to any other Peptostreptococcaceae species, reinforcing their placement within the species boundary. Core-genome recombination analyses revealed limited gene flow between cryptic and classical clades, except for Clade C7, which exhibited intermediate recombination patterns and may represent a transitional lineage. These findings expand the known diversity of *C. difficile*, provide a revised genomic framework for clade classification, and underscore the evolutionary depth and ecological breadth of cryptic lineages with potential clinical relevance.

## Introduction

*Clostridioides difficile*, a ubiquitous anaerobic spore-producing microbe, is frequently implicated in nosocomial or antimicrobial-associated infections, collectively known as *C. difficile* infections (CDI) ^1^. This environmentally persistent pathogen causes a wide range of symptoms, from mild diarrhea to severe pseudomembranous colitis ^2^. As a leading cause of healthcare-associated infections, *C. difficile* poses a significant burden on global health, with incidence rates ranging from 2 to 8 cases per 10,000 patient-days ^3,4^. Conventional treatment approaches for CDI involve antibiotics, such as vancomycin, and fidaxomicin ^5^. However, despite these interventions, approximately 15 to 35% of CDI cases experience recurrence (R-CDI) ^6-8^.

*C. difficile*’s infectious cycle begins as spores are ingested through contaminated food, water, or surfaces^9^, and germinate in the presence of a dysbiotic microbiota^10 9^. During infection, *C. difficile* can produce several toxins (toxin A or TcdA, toxin B or TcdB, and the binary toxin or CDT)^11,12^. Most *C. difficile* isolates have at least one of both, toxins TcdA and TcdB, which are responsible of the clinical manifestations of the disease^11^. By contrast, ∼ 20% of the *C. difficile* isolates encode CDT^13^, which is linked to exacerbated disease outcomes, contributing to an upsurge in morbidity and mortality^12^. These toxins are encoded in distinct chromosomal locations, with the Pathogenicity locus (PaLoc) encoding both toxins TcdA and TcdB, while the Cdt Locus (CdtLoc) carries the binary toxin (CDT)^14^. Both toxin locus are prevalent across classical clades, underscoring their clinical relevance and further extending our understanding of their distribution beyond the classical clades^15^.

The advent of whole-genome sequencing has significantly impacted our understanding of *C. difficile*’s epidemiology, composition, evolution, and differentiation. Phylogenetic analyses reveals that most *C. difficile* isolates associated with human and animal diseases belong to five main classical clades ^16^. However, several reports on phylogenetically different isolates (mostly environmental) from the five canonical clades supports the existence of several cryptic clades (C-I to C-V), which are considered unconnected genomospecies ^16-20^. The identification of these five novel ‘Cryptic’ clades (designated with Roman numerals C-I to C-V) through core-genome analysis and the average nucleotide identity (ANI) comparisons ^3, 17, 21^, reveals ANI values substantially below the 95% species demarcation threshold. This suggests that these clades are phylogenetically ancestral and diverged from the classical clades several million years ago ^15^. Importantly, isolates of cryptic clades can encode toxin genes ^16,18-20^, and recent evidence of their clinical significance^22^ highlights their infective potential. This adds a new layer of complexity to our understanding of *C. difficile* genomic diversity and intraspecies variability, underscoring the need to determine their role in human disease.

In this study, we aimed to expand our understanding of the genetic diversity and evolutionary relationships among *C. difficile* isolates from both classical and cryptic clades. We analyzed a collection of approximately 25,000 published genomes (as of 2022), supplemented with cryptic isolates sourced from Latin American (LA) countries. Through average nucleotide identity (ANI) analysis, core-genome phylogenomics, and toxin profile comparison, we provide new expanded insight into the complex genomic architecture of cryptic clades, proposed three novel cryptic clades, and highlight the potential pathogenicity of these divergent lineages.

## Materials and Methods

### Sample collection, bacterial cultivation, DNA extraction and sample screening

A total of 21 *C. difficile* strains were isolated from soil samples collected in Costa Rica (n = 20) and Brazil (n = 1) between 2003 and 2020 (Fig. S1, Table S1). Soil samples were collected from the top 10 cm of soil using sterile disposable plastic spoons at 4ºC in sterile 25 ml containers until processing. Two grams of each soil sample were incubated in Cdiff banana broth (Hardy Diagnostics) at 37°C for 72h. Cultures were then streaked onto taurocholate-cefoxitin-cycloserine-fructose agar (TCCFA) and incubated for 24 h at 37°C in a vinyl type C anaerobic chamber (Coy Laboratory Products). Single colonies were grown in BHIS supplemented with cefoxitin-cycloserine. A 20% (v/v) glycerol stock was created from this culture for future use, and DNA extracted using the Qiagen DNeasy Blood & Tissue kit, and screened with Taqman PCR assay targeting a unique *C. difficile* marker (CD630_24840), previously for detecting *C. difficile* in dog fecal samples ^*23*^. Samples testing positive by Taqman PCR were subjected to a second round of DNA extraction using the same kit for whole-genome sequencing.

### Genome collection

We obtained a dataset of 25,145 *C. difficile* genomes and associated metadata from EnteroBase, with a cutoff date of September 30, 2022 (Table S1). This comprehensive collection captures extensive strain and source diversity under a wide range of conditions. To further enhance the representation of cryptic lineages, we incorporated the 21 isolated genomes isolated in this study from Costa Rica and Brazil from suspected cryptic isolates (Table S2), thereby expanding the diversity and analytical scope of our study on cryptic clades.

### Genome assembly

Genomes were assembled, annotated, and quality-assessed using a previously described pipeline ^15^, which includes TrimGalore v0.6.5 for read trimming, SPAdes v3.6.043 for de novo assembly, Prokka v1.14.5 for annotation, and QUAST v2.3 for assembly quality evaluation ^15^. Kraken2 v2.0.8-beta was subsequently used to screen for contamination and assign taxonomic labels to both raw reads and draft assemblies.

### Multi-locus sequence typing

To identify significant genetic variations, assembled genomes were analyzed using multi-locus sequence typing (MLST) utilizing FastMLST v0.0.15 ^24^. Novel alleles and sequence types (STs) identified using this process were verified by submitting assembled contigs to the *C. difficile* PubMLST database (https://pubmlst.org/cdifficile/) ^25^, a widely used resource for molecular typing and microbial genome diversity studies.

### Average nucleotide identity

FastANI ^26^ was used to calculate Average Nucleotide Identity (ANI) across the entire dataset, providing a quantitative measure of genetic similarity between genomes. A heatmap was generated using a custom Python script to visualize ANI values. For the delimitation of cryptic clades, a 95% ANI threshold was applied, consistent with previous studies ^17^, to ensuring consistency and alignment with established species boundaries.

### Genome Subsampling Strategy

To construct a representative and non-redundant subsample of genomes for downstream analyses, we employed distinct selection strategies for classical (non-novel) clades and for cryptic or novel clades, with the goal of balancing phylogenetic diversity and assembly quality.

For the classical clades (defined as ANI>95% against *C. difficile* 630), we applied a systematic selection based on ST and assembly metrics (Table S2). Specifically, for each ST within a given classical clade, all available genome assemblies were identified and evaluated for assembly quality using the N50 metric. The genome with the highest N50 value was selected as the representative for that ST. This approach ensured that each ST was represented by a single high-quality genome, minimizing redundancy while preserving within-clade diversity. As a result, the final dataset included one genome per ST per classical clade. In contrast, for the cryptic clades (n = 125), we adopted an inclusive analyzed strategy. Due to the limited genome data available for these lineages, all associated genome assemblies were retained regardless of ST or assembly quality, to capture the full breadth of diversity within these underrepresented clades. Applying this dual strategy resulted in a curated dataset of 577 genome assemblies, representing both classical and cryptic clades with an emphasis on quality and diversity enabling robust comparative analyses.

### Core-genome genes and recombination-free phylogeny

Core-genome identification was performed using DIAMOND blastp ^27^ with a bidirectional best hit approach. Protein-coding genes were predicted using Prodigal ^28^, and *C. difficile* strain 630 was used as a reference for bidirectional best hit comparisons. Genes present in at last 99% of isolates were considered core genes. Each core gene was aligned individually using MAFFT ^29^.

To improve the accuracy of phylogenetic inference, we first accounted for the confounding effects of homologous recombination. Recombination detection and masking were performed using ClonalFrameML^30^, a tool specifically designed to identify and mask recombinant regions in bacterial genomes. Following this step, we ensured possession of a clean, recombination-free alignment. A concatenated core gene alignment, generated from the selected genome dataset, was used as input for ClonalFrameML. The tool estimated recombination events and produced a recombination-corrected alignment by masking affected regions.

The recombination-free alignment was then used to construct a maximum likelihood phylogenetic tree with IQ-tree ^31^. To evaluate the robustness of the inferred topology, we performed 1,000 ultrafast bootstrap replicates.

## RESULTS AND DISCUSSIONS

### Global distribution of *C. difficile* Clades

A prior taxonomic study of *C. difficile* analyzed over 12,000 genomes and provided a detailed insight into its taxonomic structure highlighting the existence of several cryptic clades ^15^. Given the substantial increase in available genomic data, the present study aimed to further explore the diversity landscape of *C. difficile*. We analyzed 25,145 genomes from the curated EnteroBase database (Table S1), along with 21 additional genomes from Costa Rica and Brazil sequenced in this study (Table S2). Genome assemblies varied in quality, with an average of 144 contigs per genome and an average genome size of 4.2 Mb (Table S3). Most Publicly available genomes were derived from the United Kingdom (37.2%), United States (25.8%), Germany (9.8%), Australia (2.1%), Canada (2.0%), Spain (1.8%), China (1.3%), and Japan (1.2%), while other countries total up to the remaining 7.3% (**Fig. 1A**), and the majority of genomes derived from human isolates (> 80%) (**Fig. 1B)**. The majority of genomes belonged to Clade 1 (C1, 64.3%), followed by Clade 2 (C2, 17.8%), Clade 5 (C5, 9.9%), Clade 4 (C4, 4.9%), and Clade 3 (C3, 1.8%) (**Fig. 1A**). We utilized FASTMLST to classify was based on the corresponding ST-clade associations. Among the identified STs, the most prevalent ones were ST1 (C2), ST1 (C1), and ST11 (C5) (Fig. 1D), reflecting Clade1 bias of available public database. However, FastMLST was unable to assign all STs to known clades resulting in 299 genomes lacking assignment to both ST and clade (**Figure 1C,D**), indicating limitations in the current typing schemes. Altogether, these findings suggest the presence of additional, uncharacterized diversity within the *C. difficile* population.

**Figure 1.**
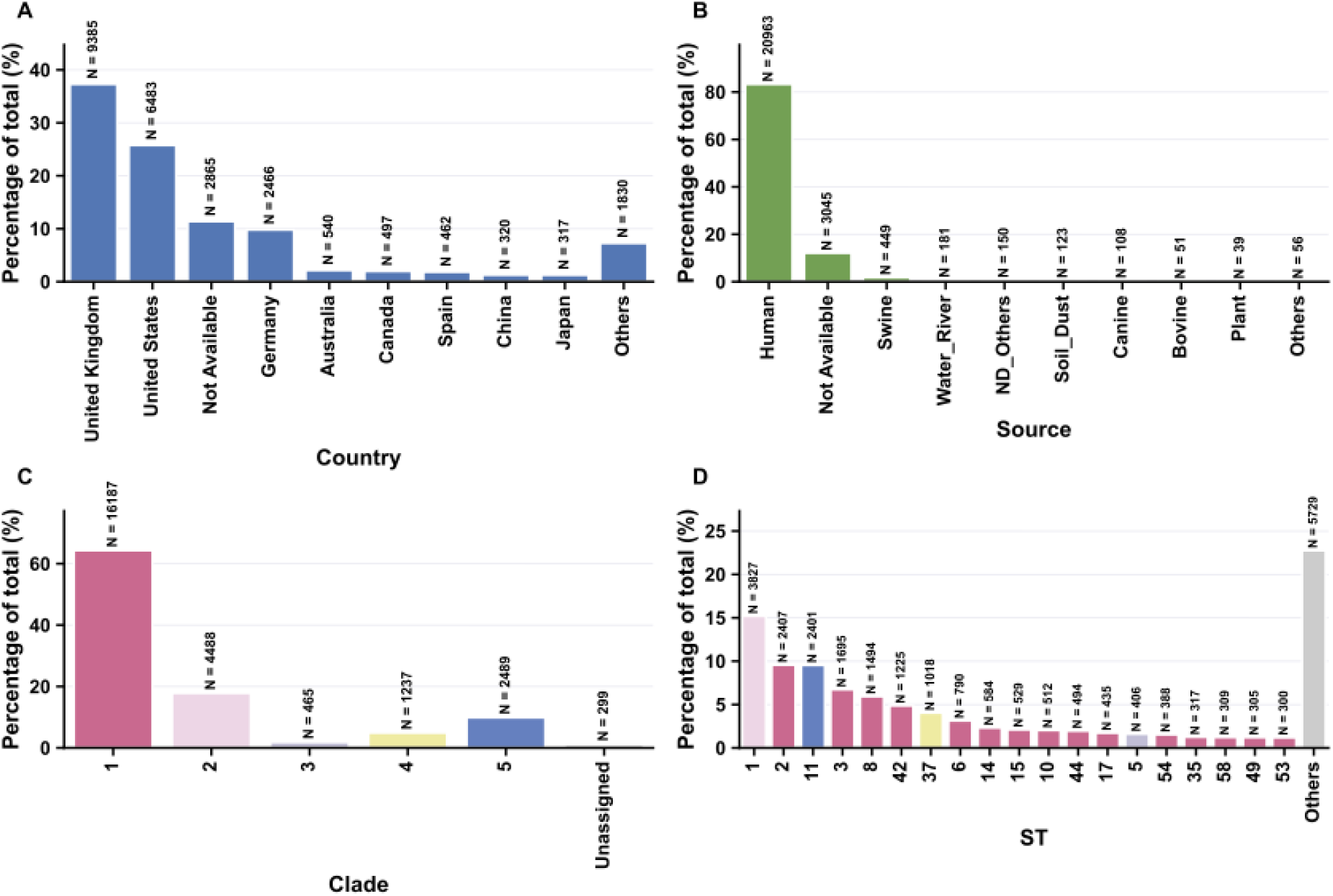
Composition of *C. difficile* genomes in the Enterobase. Snapshot obtained on 20^th^ September of 2022; 25,165 strains. A) Country and B) source of isolates. C) The proportion of genomes in Enterobase by clade. D) Top 19 most prevalent sequence types (STs) in the Enterobase colored by clade.

To visualize an updated structure of *C. difficile* global population, we constructed a rooted MLST phylogenetic tree using all 25,145 genomes (Table S3 and Figure S1). In addition, we also incorporated the known cryptic clades assignation (e.g., C-I to C-V)^17^ to our clade assignations. Our preliminary tree demonstrates the presence of all classical clades, C1 to C5, and the established cryptic clades C-I, C-II, C-III, C-IV and C-V (**Fig. S1 and Fig. 1C,D**). However, additional monophyletic nodes closely related to cryptic clades were evidenced with no clade assignment (Fig. S1). Strikingly, six additional isolates fell within the classical clade-proximity, suggesting novel lineages within these boundaries (**Fig. S1**). Altogether, these findings highlight uncharacterized diversity within the *C. difficile* population.

### Core-genome phylogeny reveals novel monophyletic classical and cryptic groups

To assign these novel lineages and explore their evolutionary relationships within *C. difficile*, we constructed a representative non-redundant subsample of the 25,145 genomes (Table S1). For this, we utilized two selection criteria, for each known ST within classical clades, we selected the highest-quality genome based on N50, resulting in 452 representative assemblies. These were then combined with 125 genomes from established cryptic clades, putative cryptic clades forming novel monophyletic groups within the cryptic boundary, and the 6 new genomes from novel classical groups. A designated outgroup, *Clostridium mangenotii* was also included due to its close but distinct evolutionary relationship with *C. difficile* (Table S4), providing a robust anchor for resolving deeper nodes and ambiguous ST-to-clade assignments, producing a final dataset of 578 genomes. The resulting tree robustly resolved the classical clades C1 to C5 with strong bootstrap support (>90 for all split clade nodes) (Fig. 2A). Strikingly, two novel STs were identified, ST284, represented by five genomes, was positioned at the boundary of C1 and C2, while ST881, represented by a single genome (CLO_CA5670AA), was positioned at the boundaries of the known classical clades (Fig. 2A). Both, ST284 (n = 5) and ST881 (n = 1), are proposed as novel clades within the classical boundaries and named C6 and C7, respectively (Fig. 2A). Additionally, two novel cryptic clades were also identified, which we designated C-VI and C-VII, as well as a sub-lineage within cryptic C-III clade (e.g., C-IIIa and C-IIIb) (Fig. 2A,B).

**Figure 2.**
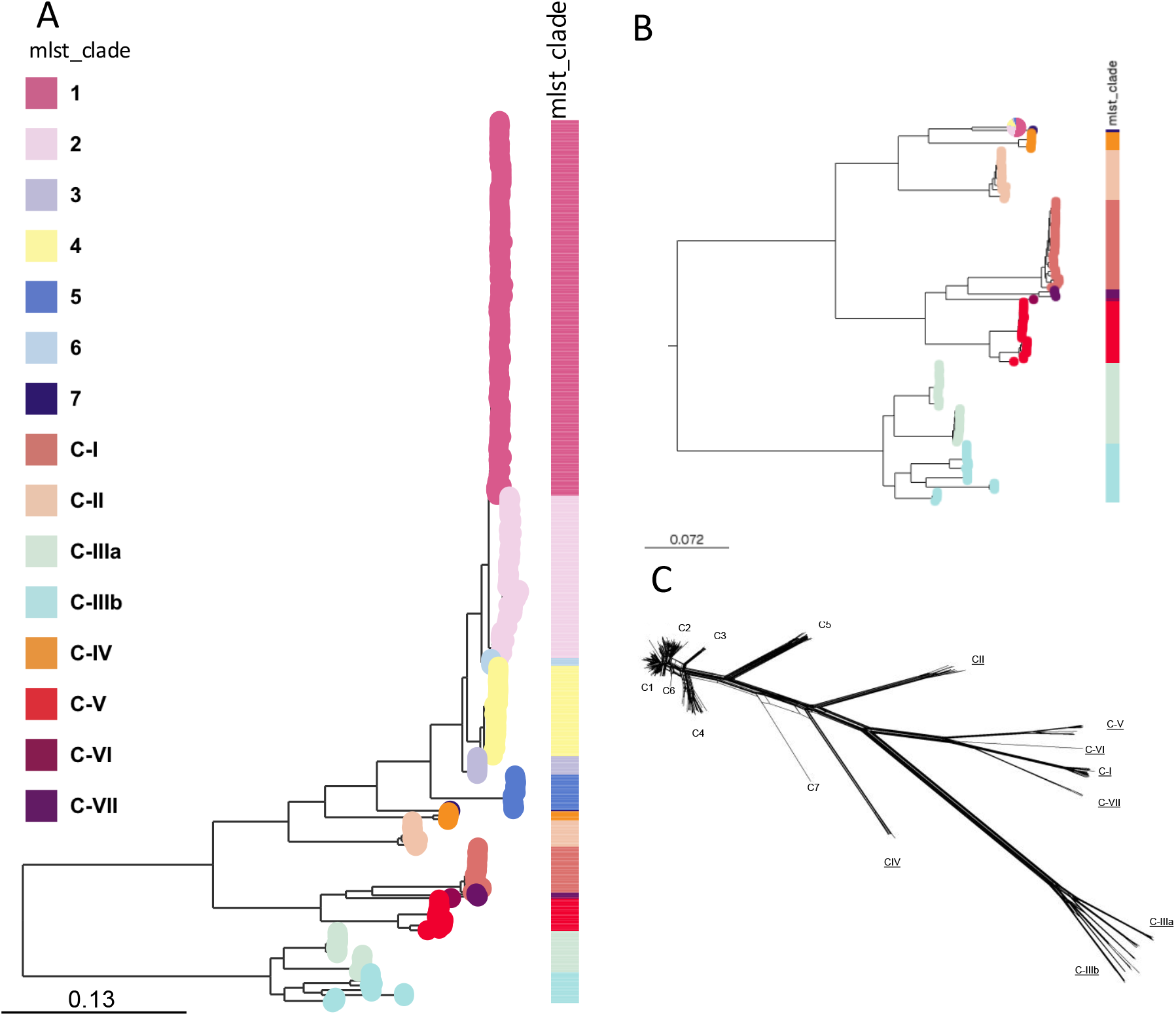
Phylogenetic Relationships and of *C. difficile* Isolates. (A) The recombination-free phylogenetic tree showcases the relationships among various C. difficile isolates. The tree incorporates *Clostridium mangenotii* as an outgroup to root the tree and provide context. The color-coded clades represent distinct evolutionary lineages, with the classical clades C1-C5 known prior to this study and the newly identified clades C6 and C7 highlighted. The core genes were identified and aligned, ensuring no recombination events to maintain accuracy. All split nodes dividing between clades have Bootstrap values > 90. (B) Recombination network analysis depicting gene flow dynamics across clades. While Clades C1, C2, C3, C4, and C6 are interwoven with dense connections indicating substantial gene exchanges, Clade C5 stands in contrast, reflecting limited gene flow. The standout observation is the unique positioning of Clade C7, which seems to act as an intermediary, establishing bridges between classical and cryptic clades. This suggests C7’s potential role as a genetic hub, enabling the exchange of genetic material between these groups. Patterns of reticulation between C7, classical clades, and C-IV underscore this hypothesis. In contrast, cryptic clades like C-I, C-II, C-IV, C-V, C-VI, and C-VII largely remain in isolation, with limited reticulations, except for C-IIIa and C-IIIb which show pronounced gene flow at their divergence point.

To assess gene flow between clades, we conducted a recombination network analysis using the core-genome alignment (Fig. 2C). We observed that clades C1, C2, C3, C4, and the proposed C6 displayed highest levels of reticulation, indicative of frequent recombination events. In contrast, Clade C5 exhibited lower levels of gene flow with both classical and cryptic clades, suggesting a more genetically isolated lineage. However, clade C7, although not considered cryptic due to its ANI > 95% with strain 630, appeared phylogenetically close to C5 and exhibited an intermediate recombination profile (Fig. 2C). Classical clade C7 shows evidence of recombination with both, classical and cryptic clade, suggesting that C7 may act as a transitional lineage facilitating *gene exchange across the boundaries between classical and cryptic clades* (Fig. 2C). Most cryptic clades C-I, C-II, C-IV, C-V, C-VI, and C-VII show limited evidence of recombination, suggesting limited gene flow among them and possible evolutionary isolation (Fig. 2C). An exception was observed in cryptic clade C-III, where the sub-lineages C-IIIa and C-IIIb showed evidence of extensive gene flow early in their divergence. Collectively, these findings highlight the role of Clade C7 as a potential genetic bridge, facilitating horizontal gene transfer between classical and cryptic clades.

### Average nucleotide-identity (ANI) analysis reveals novel classical and cryptic clades in *C. difficile*

To quantitatively delineate these novel monophyletic groups and address their genomic similarities, we utilized whole-genome ANI analyses. Whole-genome values were determined for the final set of 577 genomes (without the outgroup *C. mangenotti*) using FastANI. To more accurately classify novel clades of *C. difficile* isolates within our dataset, we reassigned each isolate to a unique ST and designated as cryptic if its ANI value was < 95% relative to reference strain 630. ANI analysis revealed a large cluster of genomes corresponding to the classical *C. difficile* clades (e.g., C1 to C5) with ANI values higher than 95% (Fig. 3A), along with several smaller, distinct clusters. However, given the low number of genomes of the proposed novel clades 6 and 7, they are undetectable in ANI heat map analyses in Fig. 3A. Notably, upon closer inspection of these smaller clusters, highlighted within the white segmented box (Fig. 3A,B), we identified all five previously described cryptic clades: C-I (n = 30), C-II (n = 17), C-IV (n = 6), and C-V (n = 21), as described in prior studies^17^. Importantly, upon closer examination of the highlighted area in Fig. 3A, we were able to detect the two novel cryptic clades: C-VI (n = 1) and C-VII (n = 3) (Fig. 3B). Moreover, within the original C-III clade, the pairwise-ANI heatmap revealed two distinct blocks of genomes separated by ANI values below 95%. This violates the empirical intra-clade coherence criterion, namely, that all members of a given clade should share ≥ 95% within each other, a rule consistently upheld by all other clades in our dataset (Supplementary Figure S1). Interestingly, partitioning the previously described Clade C-III (n=27) into C-IIIa^32^, and a newly defined C-IIIb (n = 20), restores that coherence in this clade, in that all intra-group comparisons remain ≥ 95 % (Fig. 3.

Closer examination of the ANI comparisons of the values confirms that the known classical clades, Clades C1 to C4, cluster together within an ANI value of 98% (Fig. S2A,B,C,D), whereas classical C5 cluster with ANI-values of 96% against Clade C1 to 4 (Fig. S2E). Notably, cryptic clades C-I, C-II, C-IV and C-V all had values between 89 to 95% (Fig. S2F,G,H,I,J), consistent with the notion of genomospecies and with their prior classification as cryptic lineages ^16,17^. Notably, among the new classical clades, C6 shared ANI-values between 98.1 to 98.7 with classical clades C1 to C4 and 96.1 with C5 (Fig. 4A), confirming that C6 is closely related to classical clades C1 to C4. By contrast, C7 had ANI-values in the range of 95.0 to 95.8 with classical clades C1 through C6 (Fig. 4B), whereas ANI-values ranged from 89.1 to 94.1% with cryptic clades (Fig. 4B), which agrees with C7 being at the interface between both classical and cryptic clades.

**Figure 3.**
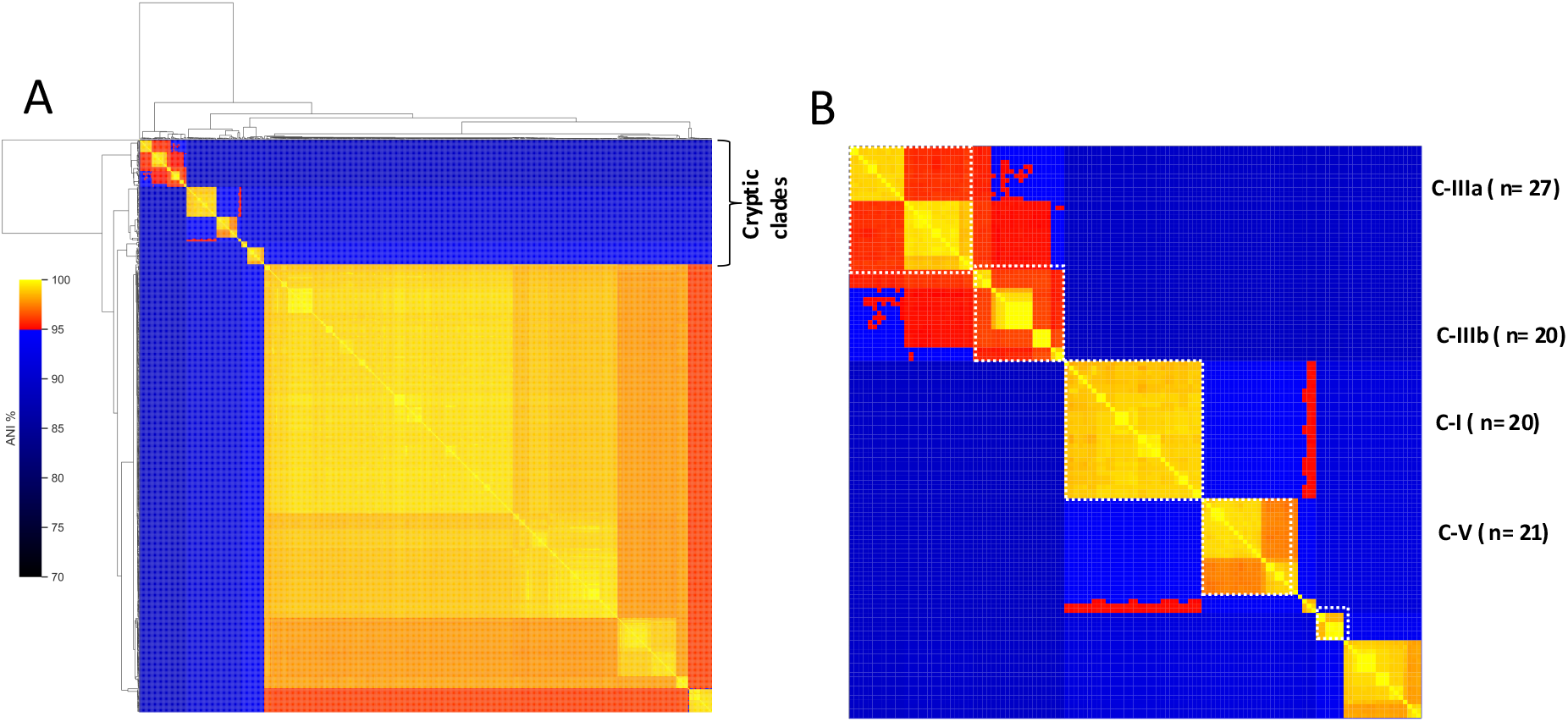
Average Nucleotide Identity *of Novel Cryptic* C. difficile genomes. (A) Heatmap showcasing the Average Nucleotide Identity (ANI) to highlight genetic similarity among C. difficile genomes. The color gradient, from blue to red, indicates ANI percentage; values nearing 100% signal increased genetic resemblance. An ANI threshold of 95% was utilized to identify cryptic clades. Genomes with ANI below this mark are encompassed within the white segmented box, demarcating them from the broader C. difficile genomic landscape. (B) A focused view of the cryptic clades extracted from panel (A). While classical clades are illustrated as C-I through C-V, this research unearthed two novel cryptic clades, denoted as C-VI and C-VII. Additionally, a bifurcation within clade C-III led to its division into C-IIIa (previously recognized as C-III) and C-IIIb.

**Figure 4.**
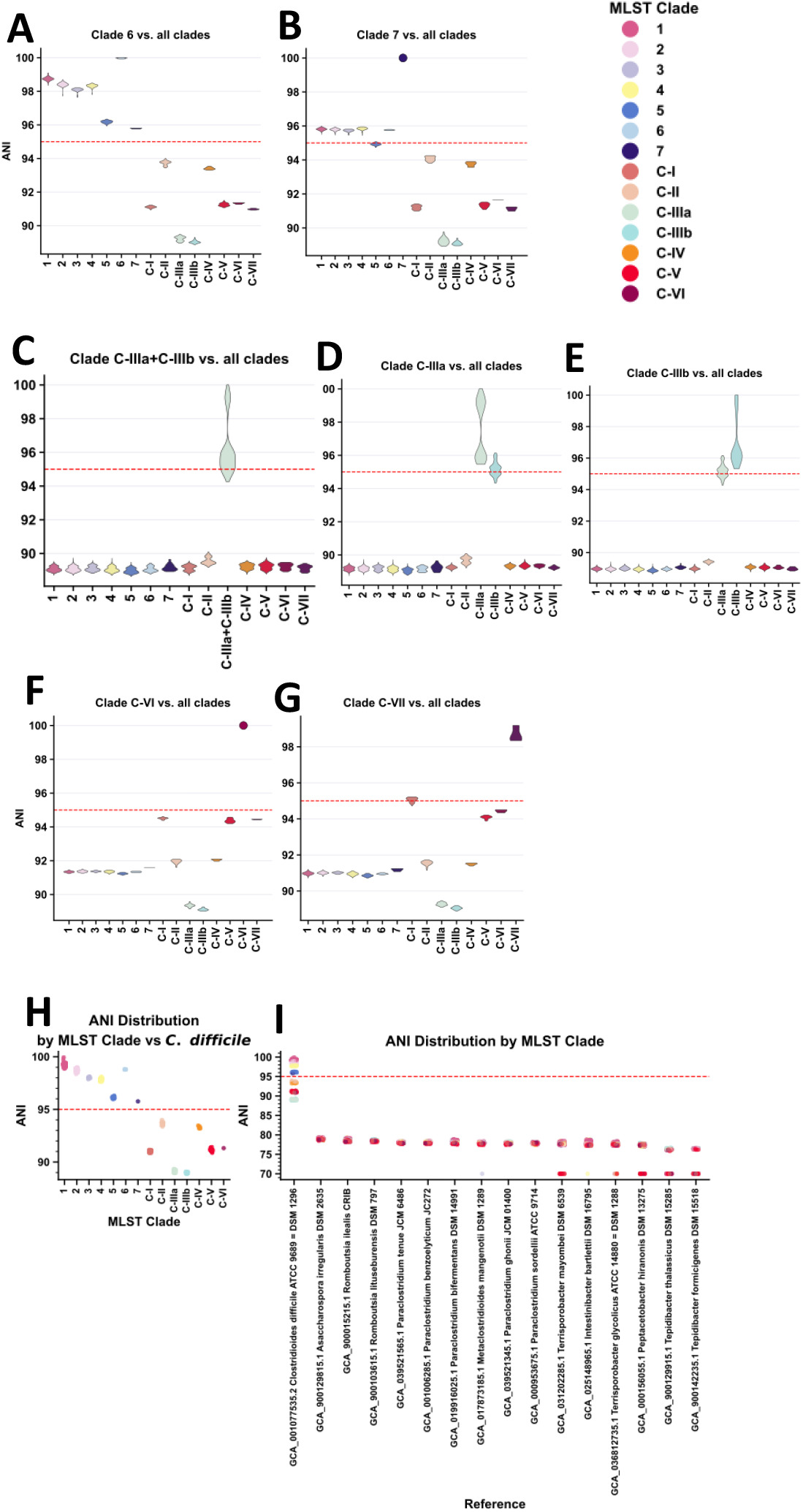
Analysis of ANI distributions. A) Violin plot of ANI values between pairs of 571 genomes grouped by clade. Notably, all defined clades exhibited ANI values >95%, a threshold traditionally used for species delimitation, reinforcing the cohesiveness of these groups within the species. This high ANI value within clades indicates that they function as cohesive genetic units. A boxplot is shown inside the violins where the white dot indicates the median of the data B) Scatterplot plot of ANI values between pairs of 571 genomes compared against the reference genome of all the Peptostreptococcaceae family. A horizontal red line species/lineage boundary of 95 % ANI.

FAST-ANI analysis of cryptic clade C-III confirmed that this clade seems to be at early stages of bifurcation (Fig. 4C). Dividing C-III into C-IIIa and C-IIIb as stated above reveals demonstrates that ANI-value between with C-IIIa and C-IIIb is 95.2%, supporting the notion that they are at the border line of being same species (Fig. 5D and 5E). Notably, both of these cryptic clades shared ANI-values with classical and cryptic clades in the range of 89 to 89.5% (Fig. 5D and 5E), further supporting the notion of emergence into a new genomospecies. Notably, the putative cryptic clade C-VI and C-VII, had ANI-values with other cryptic clades from 89.1 to 94.5% and 89.0 to 95.1, respectively (Fig. 5F and 5G), supporting these monophyletic groups as novel cryptic clades.

Figure 4H provides a zoomed-in view of ANI values specifically between each MLST clade and the reference *C. difficile* genome. As shown, classical clades and the newly proposed C6 and C7 consistently exceed the 95% ANI threshold, while cryptic clades range from just under 95% down to approximately 90% (Fig. 5H). This zoomed perspective highlights the considerable genomic divergence present among cryptic clades, yet also reinforces that all of them are more closely related to *C. difficile* than to any other species in the family. The gradient of ANI values across cryptic clades may reflect heterogeneous evolutionary histories, shaped by niche specialization and varying selective pressures.

To place the novel classical and cryptic clades identified in this study into a broader taxonomic and evolutionary context, we evaluated their genomic relatedness to other members of the *Peptostreptococcaceae* family (Fig. 5I). This comparison aimed to determine whether these lineages represent highly divergent branches within *C. difficile* or should instead be considered distinct species.

Although cryptic clades exhibit ANI values below 95% when compared to the reference *C. difficile* strain a threshold commonly used for species delimitation (Fig. 5I), they remain clearly more similar to *C. difficile* than to any other species within the family, displaying ANI values consistently below 85% with non-*C. difficile* taxa (Figure 6A). This finding supports the interpretation that, despite their deep divergence, cryptic clades fall within the species boundaries of *C. difficile*.

Rather than constituting separate species, these lineages likely represent evolutionarily ancient branches that have maintained a stable *C. difficile* core genome while accumulating extensive genomic divergence. Such a pattern, high intra-species divergence coupled with inter-species separation, is not uncommon in bacterial species with long evolutionary histories. It reflects a dynamic evolutionary landscape shaped by the combined effects of vertical inheritance and episodic horizontal gene transfer. This balance of conservation and diversification supports the hypothesis that cryptic clades are highly specialized genomic lineages that evolved within the boundaries of *C. difficile*, expanding our understanding of its genomic plasticity and taxonomic complexity.

## Conclusion

In conclusion, our study offers valuable insights into the genetic diversity and evolutionary history of *C. difficile* clades by confirming previously reported cryptic clades, identifying two additional cryptic clades (C-VI and C-VII), and suggesting a potential subdivision of clade C-III into C-IIIa and C-IIIb. These findings emphasize the importance of continued research to further understand the epidemiology, pathogenicity, and clinical management implications of *C. difficile*’s complex and diverse clades.

## Supplementary Figures

**Figure S1**. MLST Diversity and Distribution of *C. difficile* Clades and Sequence Types (STs). (A) Distribution of *C. difficile* clades based on 25,165 genomes (25,144 from Enterobase and 21 cryptic genomes Snapshot taken September 30, 2022 from Enterobase). Clade C1 predominates, making up 64.3% of all genomes analyzed, followed by Clades C2, C5, C4, and C3 at 17.8%, 9.9%, 4.9%, and 1.8% respectively. Notably, 299 genomes are unassigned to any clade. (B) Prevalence of identified STs with more than 100 isolates. ST1 from both Clade C2 and C1 are among the most common, followed closely by ST11 from Clade C5. The study suggests significant underlying diversity within *C. difficile* and its cryptic clades.

**Figure S2. Analysis of ANI distributions**. A-G) Violin plot of ANI values between pairs of 571 genomes grouped by clade. Notably, all defined clades exhibited ANI values >95%, a threshold traditionally used for species delimitation, reinforcing the cohesiveness of these groups within the species. This high ANI value within clades indicates that they function as cohesive genetic units. A boxplot is shown inside the violins where the white dot indicates the median of the data. I) Scatterplot plot of ANI values between pairs of 571 genomes compared against the reference genome of all the Peptostreptococcaceae family. A horizontal red line species/lineage boundary of 95 % ANI

