## Supplementary figures and images for "Identification of Novel Cryptic and Classical Clades in *Clostridioides difficile*"

### Figure S1,2

#### Figure S1

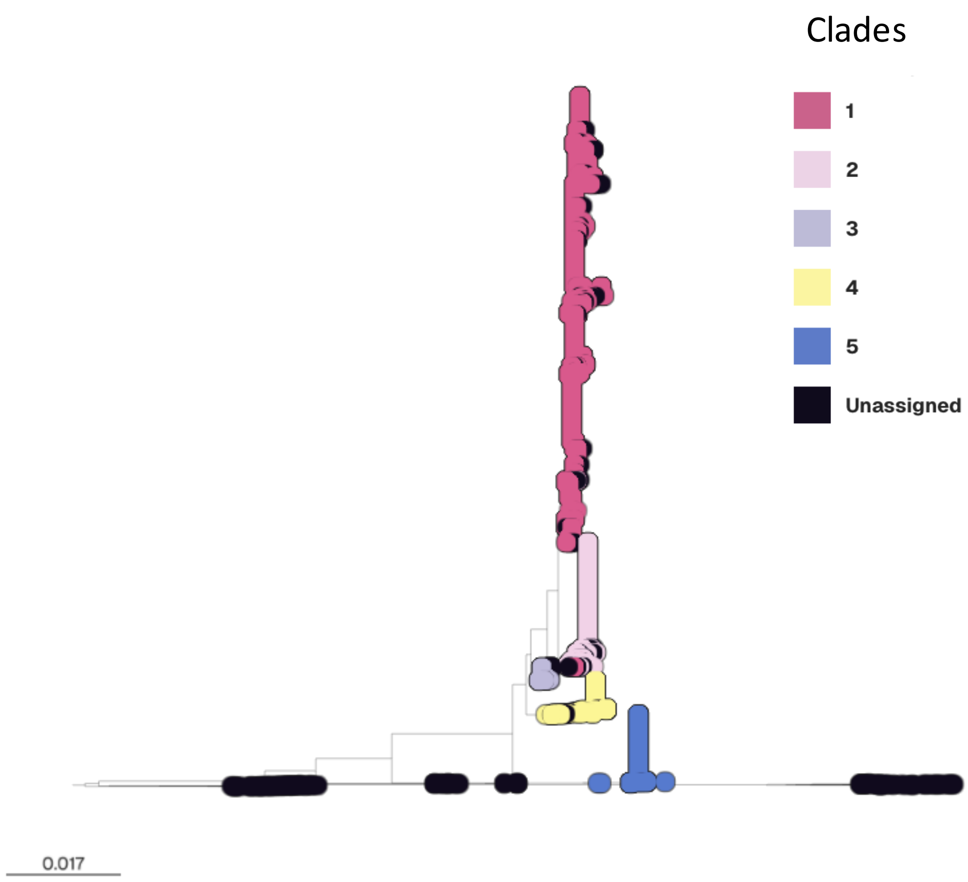

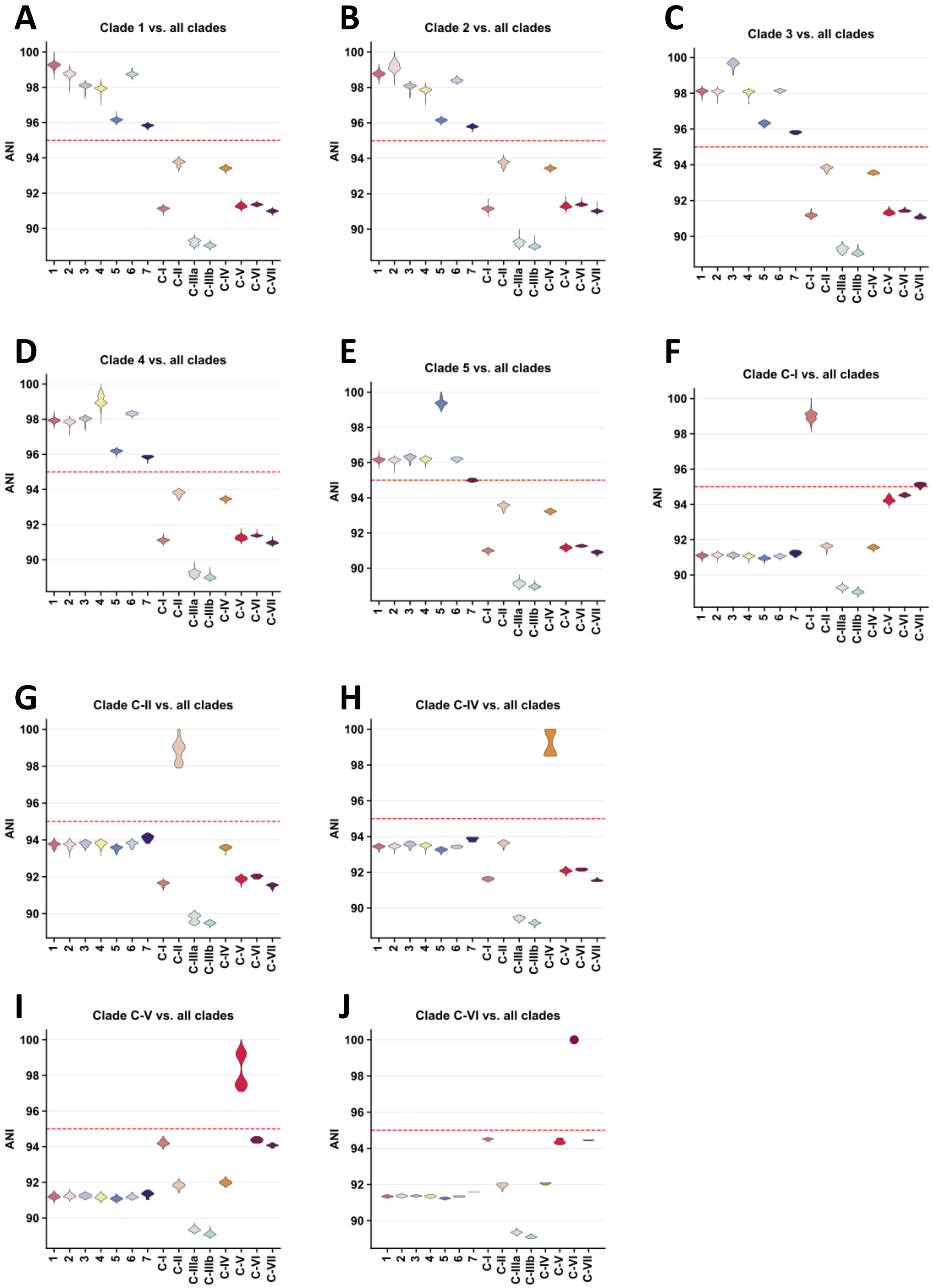
